# Unsupervised learning reveals landscape of local structural motifs across protein classes

**DOI:** 10.1101/2023.12.04.569990

**Authors:** Alexander Derry, Russ B. Altman

## Abstract

Proteins are known to share similarities in local regions of 3D structure even across disparate global folds. Such correspondences can help to shed light on functional relationships between proteins and identify conserved local structural features that lead to function. Self-supervised deep learning on large protein structure datasets has produced high-fidelity representations of local structural microenvironments, enabling comparison of local structure and function at scale. In this work, we leverage these representations to cluster over 15 million environments in the Protein Data Bank, resulting in the creation of a “lexicon” of local 3D motifs which form the building blocks of all known protein structures. We characterize these motifs and demonstrate that they provide valuable information for modeling structure and function at all scales of protein analysis, from full protein chains to binding pockets to individual amino acids. We devise a new protein representation based solely on its constituent local motifs and show that this representation enables state-of-the-art performance on protein structure search and model quality assessment. We then show that this approach enables accurate prediction of drug off-target interactions by modeling the similarity between local binding pockets. Finally, we identify structural motifs associated with pathogenic variants in the human proteome by leveraging the predicted structures in the AlphaFold structure database.

## 1 Introduction

Systematic classification of protein structures is important for understanding evolutionary relationships between proteins and the principles of how structure leads to function. As databases of high-quality protein structures expand rapidly, computational methods for automatically characterizing protein structure and function are necessary to fully realize the potential of the wealth of new data. Currently, protein structures are typically classified at the domain level (continuous regions of about 50–200 amino acids) using databases such as SCOP^1, 2^ and CATH^3^, which assign domains to a hierarchy of unique three-dimensional folds. Since large proteins are often made up of multiple domains, identifying conserved structural conformations at the domain level has proved very useful for discovering distant evolutionary relationships.

However, the idea that structures can be uniquely classified into a discrete space of conformations is limiting and can obscure the full range of structural and functional relationships between proteins^4–7^. A single structural family can contain proteins with many different functions (e.g. enolases, TIM (*α*/*β*)8 barrels), and some functions can be shared across proteins with entirely different folds (e.g. ATP/NAD/FAD binding)^7, 8^. Additionally, protein function is a nebulous concept that can be defined at many different scales, and while some aspects of function can be predicted well by domain-level classifications (e.g. global enzymatic activity), others require a more granular approach (e.g. ligand binding, post-translational modifications). Therefore, approaches based on identifying local structural features shared across fold space have been proposed as a complementary approach to domain-based classification schemes^5^.

Fragments, or local regions of 5–20 consecutive amino acids, are a popular choice of sub-domain representation which led to many new methods for functional characterization as well as advancements in protein structure prediction and de novo design^5, 9–12^. This success was due to the fact that local fragments are observed to cluster into a limited number of conserved “building blocks” which can be assembled in various ways to produce the wide diversity of folds observed across the proteome^13^. However, fragment-based approaches still require that a motif be represented by a continuous polypeptide chain, limiting their ability to capture motifs that involve amino acids far apart in sequence space (e.g. the catalytic triad in serine proteases).

Representations based on 3D microenvironments address this issue by directly modeling the configuration of atoms in a local region of 3D space independently from their position in the protein chain. Common approaches have included FFFs^14, 15^, reverse templating^16^, and PDBSpheres^17^, which use atoms from a small number of key residues to define motifs and threading or local structure alignment to match motifs to new structures. FEATURE^18, 19^ took a different approach by building a vector representation of the microenvironment around a site using a set of hand-crafted features. Vector representations are more efficient to compare than 3D structural alignments and have enabled discovery of functional similarities across diverse families^20, 21^, selectivity and off-target profiling of kinases^22, 23^, and improved modeling of protein-ligand binding^24, 25^. Unsupervised clustering of FEATURE representations across proteins has been shown to recapitulate known sequence motifs^26^ and enable discovery of novel functional patterns^20^, but this analysis has been limited to small subsets of the Protein Data Bank (PDB) and specific types of sites (e.g. disulfide bonds). Additionally, FEATURE vectors do not account for full atomic configuration, are difficult to use computationally due to a mix of categorical and ordinal features, and have limited information content due to their reliance on hand-crafted features.

The advent of deep representation learning on biomolecular structure has been transformative, enabling significant improvements over previous methods in structure prediction^27, 28^, protein design^29–31^, and many other tasks^32–34^. Recently, Foldseek^35^ introduced a variational autoencoder trained to discretize local backbone conformations into a 20-letter structural alphabet which has been effective at increasing the speed and sensitivity of structural database search, and this alphabet has shown early promise in improving protein language modeling^36, 37^. However, an alphabet of 20 letters is too coarse to capture the full variation in microenvironments that we expect to see across the proteome. Additionally, representations based on local backbone configuration are appropriate for representing general fold but modeling many functional sites requires consideration of the side chain as well as backbone atoms. We recently published COLLAPSE, a self-supervised learning method which leverages evolutionary conservation between protein families to produce learned embeddings of local microenvironments in protein structure at atomic resolution, a major improvement over previous approaches to local structure representation^38^. In the COLLAPSE embedding space, sites with similar structural and functional roles have similar representations regardless of their global sequence or fold family, enabling improvements in the classification, search, and annotation of functional sites^38, 39^.

In this work, we investigate whether the geometry of the COLLAPSE embedding space can be used to characterize the relationships between local structural sites at proteome scale. To this end, we leverage COLLAPSE embeddings to cluster over 15 million protein microenvironments in the PDB. By associating each cluster with biochemical and functional metadata, we can begin to more fully understand the building blocks which contribute to protein structure and function. We show that these clusters form a “lexicon” of structural sites that can be used to effectively model proteins at a site level, and that the cluster frequencies within a protein or protein region (e.g. binding site) can be used to create powerful representations for structural and functional analysis. Finally, by mapping clinical variant annotations onto the AlphaFold-predicted structures for the human proteome, we identify clusters which are enriched for pathogenic variation, suggesting the critical role of certain structural motifs for protein function.

## 2 Methods

### 2.1 Data processing and embedding

To account for all solved protein structures in our clustering, we used a subset of 79,324 chains from the PDB with redundancy reduction at 100% sequence identity, resolution better than 3.5 Å, R-value less than 1.0, and length between 40 and 1200 amino acids. These chains were retrieved using the PISCES server^40^. We then embedded the environment around every standard amino acid in each chain using COLLAPSE. Each environment is comprised of all atoms within a sphere of 10 Å radius centered around the centroid of the functional side-chain atoms of the central residue, and the vector output by COLLAPSE represents a learned mapping of the salient structural and functional features of this environment into a 512-dimensional Euclidean space^38^. Non-protein heteroatoms (e.g. bound ligands) were removed so they did not bias the clustering. The resulting training dataset consisted of 15,030,977 microenvironment embeddings, each mapping to a unique PDB chain and residue identifier.

### 2.2 Clustering procedure and evaluation

For computational efficiency, we clustered the embeddings using mini-batched *k*-means clustering with a batch size of 4096 parallelized over 8 CPU cores. Before clustering, we L2-normalized all embeddings to enable better comparison between points in high dimensions. We varied the number of clusters *k* in order to find an optimal value for representing the variation in structural sites, testing values of *k ∈ {*20, 100, 1000, 5000, 10000, 25000, 50000, 100000*}*. For each value of *k*, we used three random initializations and selected the best run using the entropy of SCOP families within each cluster (see Section 2.3) and total within-cluster sum of squares 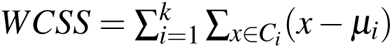, which measures the intrinsic compactness of the clustering in the absence of external labels. We chose a value of *k* = 50, 000 for all future analysis by visual inspection (Fig. S1). Note that the increase in WCSS from *k* = 50, 000 to *k* = 100, 000 is likely due to variance caused by the low number of initializations (required to complete clustering with high *k* on such a large dataset in less than 24 hours).

### 2.3 Cluster assignment and characterization over entire PDB

To create the full cluster database, we embedded the environment around every protein in the PDB (171,679 structures) using the procedure described in Section 2.1, resulting in 104,863,360 unique embeddings. We then mapped each embedding to a cluster using the clustering model trained in 2.2. Using these mappings, we identified the set of all residues belonging to each cluster. Then, we extracted the following biochemical features of each residue from its PDB entry and linked external databases.

#### Characterization from intrinsic structure

- **Central residue identity.** We computed counts of the central amino acid type for each environment in the cluster. We then computed the entropy *S* of the distribution over the set of standard amino acids *{AA}*, a measure of how conserved the central residue is:

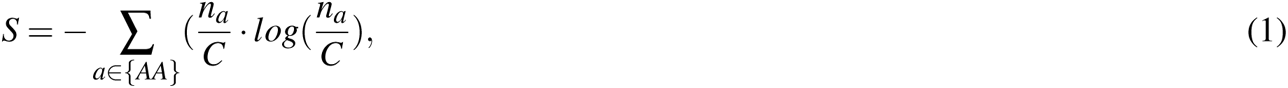

where *n_a_*is the count of amino acid *a* and *C* = ∑*_a∈{AA}_ n_a_*. We assess the substitutability of different residues in the same structural environment by measuring the pairwise correlation between the frequency distributions of each amino acid across clusters. Specifically, if we consider a *k ×* 20 frequency matrix where entry (*i, j*) is the fraction of environments in cluster *i* that are centered around amino acid *j*, we compute the Pearson correlation between all 190 possible pairs of columns to produce a 20 *×* 20 correlation matrix.
- **B-factor.** We extracted the cystallographic B-factor for the central residue from the PDB file. This quantity is related to the level of order in the structure at that position.

#### Characterization from external databases

- **SCOP classification.** We computed the structural domain classification for each environment using the residue-level mappings for each domain defined in SCOP 2.08^1^. Entropy over SCOP families was computed the same way as entropy over residues (Eq. 1).
- **Ligand binding.** We downloaded data for all non-protein ligands in the PDB from the BioLIP database^41^. We found the clusters corresponding to each residue annotated as in contact with a ligand and created a mapping from each cluster to the set of all ligands known to be in contact with it. For enzymes, we also record all residues which are annotated as catalytic.

### 2.4 Construction of protein fingerprints

First, we embed the environment around every residue and compute the corresponding clusters, so that a protein with *n* residues contains *n* clusters. We then devised a protein fingerprint representation based on the frequency of each cluster. Specifically, we create a *k*-dimensional vector *F* in which element *i* represents the frequency of cluster *i* in the protein *p* as follows:

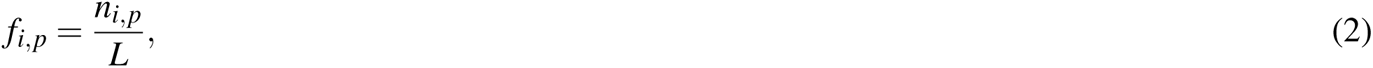

where *n_i,p_*is the raw count and *L* is the length of the protein. To account for differences in cluster specificity, we normalize each element by the inverse frequency of proteins containing cluster *i* in the entire dataset *P*. Therefore, each element *F_i_* is computed as follows:

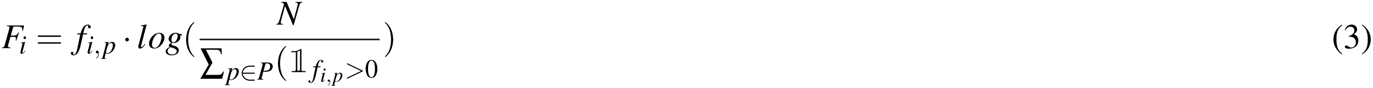

The dataset *P* over which this normalization is performed could be task-specific or based on cluster frequencies over a large protein dataset such as the PDB. We use task-specific normalization for all experiments.

#### Fold search on SCOP40 dataset

We evaluated the ability of our cluster-based fingerprints to perform protein structure search using the same dataset used to evaluate Foldseek, which contains 11,211 proteins derived from SCOP 2.01^35^. We removed all proteins that do not share a family-level classification with any other proteins in the dataset, resulting in 8,975 proteins. We then performed an all-vs-all comparison of these structures by computing the pairwise cosine similarity of their cluster fingerprints. For each query protein, we sorted all other proteins by similarity and computed the precision 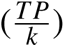 of the top *k* proteins in the ranked list. We computed the number of true positives *TP* at three levels of specificity based on the SCOP hierarchy: family (most specific), superfamily, and fold (least specific). We compared our results to an all-vs-all Foldseek search using the `easy-search` protocol with standard parameters (`--threads 32 -s 9.5 --max-seqs 2000 -e 10`).

#### Protein structure ranking benchmark on ATOM3D dataset

We used the Protein Structure Ranking (PSR) benchmark dataset from the ATOM3D project^32^ to evaluate the ability of our protein fingerprints to assess the quality of a predicted protein structure, a task that is also known as model quality assessment (MQA) or estimation of model accuracy (EMA). The dataset was derived from the Critical Assessment of Structure Prediction (CASP) challenges^42^ and is made up of the experimental structures (“targets”) and corresponding computational structure predictions (“decoys”) submitted to the competition in each year. The dataset is split into train, validation, and test sets based on the year of the competition to mirror a realistic prediction scenario. We trained a simple linear prediction head (fully-connected layers followed by ReLU activation and dropout) on top of the cluster-based fingerprints to predict GDT-TS, a numerical metric representing the overall similarity of the decoy structure to its target. To optimize this predictor, we perform a gridsearch over the following set of hyperparameters: learning rate *∈ {*1 *×* 10*^−^*^5^, 1 *×* 10*^−^*^4^, 1 *×* 10*^−^*^3^*}*, hidden size *∈ {*1024, 2048, 4096*}*, number of hidden layers *∈ {*1, 2, 3*}*, dropout rate *∈ {*0.25, 0.5, 0.75*}*, weight decay *∈ {*0.001, 0.01, 0.1*}*. Each model was trained using the Adam optimizer^43^ with standard parameters for up to 100 epochs, with early stopping used to select the best model according to mean-squared-error loss on the validation set. The best model had learning rate 1 *×* 10*^−^*^5^, 2 hidden layers with dimension 2048, dropout of 0.5, and weight decay of 0.001. We report mean and standard deviation of this model on the held-out test set over three random initializations, and compare to the benchmark results reported in Townshend *et al.*^32^.

### 2.5 Kinase pocket similarity and selectivity profiling

We obtained kinase data from the Kinase-Ligand Interaction Fingerprints and Structures (KLIFS) database^44^, which defines the binding pocket of every kinase in the PDB using a standardized set of 85 residues. We considered only pockets with a quality score of at least 6 (out of 10) and resolution at least 3.5 Å. This resulted in a dataset of 7206 structures for 283 unique kinases, each of which belonged to one of 95 kinase families and 9 kinase groups: AGC, CAMK, CK1, CMGC, STE, TK, TKL, Atypical, and Other. We embedded the environment around every pocket residue (without considering the bound ligand) and computed its cluster, resulting in 85 clusters for each pocket. We then created fingerprints (Section 2.4) for each pocket, removing cluster indices which were not represented in the dataset (i.e. zero across all examples).

We quantified the ability of the pocket fingerprints to capture kinase families using a silhouette score, which reflects how similar pockets within a kinase family are relative to those belonging to different families. We computed a silhouette coefficient *S* for each pocket 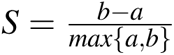, where *a* is the mean cosine distance between the pocket and all other pockets in the same family, and *b* is the mean cosine distance between the pocket and all pockets in the next closest family. We compared the performance of our fingerprints to the KLIFS interaction fingerprint (IFP), a bit vector representing the protein-ligand interactions present in the structure of a particular complex, including hydrogen bonding, ionic bonding, and hydrophobic contacts^45^. We used a Jaccard distance metric for comparing KLIFS IFPs. Statistical significance for the difference between silhouette scores for each family was assessed using a Bonferroni-corrected p-value cutoff of *p <* 0.05 computed using the Wilcoxon signed-rank test. Only families with 10 or more examples were considered for significance testing.

To assess how well fingerprint similarity could capture kinase selectivity to small molecule inhibitors, we used binding data published by Karaman *et al.*^46^, which measured the interactions between 38 inhibitors and 317 kinases. We focused on 9 kinase inhibitors with clinical relevance and measured affinity to at least 50 targets to enable accurate evaluation (Table S1. We separately computed the selectivity profile of each inhibitor with respect to all of its primary targets (as specified by Karaman *et al.*). Specifically, we computed the cosine similarity between the target pocket and all other pockets in the dataset, resulting in a list of predicted off-target kinase interactions ranked by similarity to the primary target. For kinases with multiple structures, we computed all pairwise similarities and chose the target-kinase pair with the highest similarity. We evaluated this ranking relative to the experimental data, which we binarized to define positive binding interactions using a cutoff of 100 nM. Our primary evaluation metric was the average precision (AP) relative to positive binding interactions over all thresholds of pocket similarity score. We compared to the same procedure applied to KLIFS IFPs using Jaccard similarity, since this is the primary metric used for pocket similarity searching in the KLIFS database^44^.

### 2.6 Human clinical variant analysis

We obtained variant annotations for all missense variants in human UniprotKB/SwissProt^47^ entries from the Humsavar database (http://www.uniprot.org/docs/humsavar; Release 2023_04)^48^. This database contains 82,412 variants, of which 32,627 are annotated as “pathogenic” or “likely pathogenic”, 39,654 are “benign” or “likely benign”, and 10,131 are of unknown significance. To ensure full coverage of the proteome and accurate mapping of residue positions, we used structures predicted by AlphaFold2 (https://www.alphafold.ebi.ac.uk/download)^27, 49^. We embedded these structures using the same procedure described above and computed the cluster for all residue microenvironments using the pre-trained clustering model, considering only positions with pLDDT *>* 70. Then, we mapped all variant annotations in Humsavar to their corresponding cluster using their Uniprot residue index. To identify clusters which are enriched for pathogenic mutations, we performed a hypergeometric enrichment test in each cluster and applied a Bonferroni correction to account for multiple hypotheses.

## 3 Results

### 3.1 Characterization of clusters across the PDB

By fitting clustering models on the entire non-redundant PDB, we learned a deterministic mapping from any local microenvironment in 3D space to a discrete element in a “lexicon” of 50,000 structural motifs (see Methods 2.2). We applied our trained clustering model to assign a cluster to every residue in the entire PDB (104,863,360 environments). The resulting clusters vary in size from 1 to 187,002, with over 90% containing less than 4,000 environments (Fig. 1A). The very large clusters in the right tail of the distribution appear to be made up largely of small residues (e.g. glycine, alanine, serine) and are predominantly found in nuclear magnetic resonance (NMR) and low-resolution cryo-electron microscopy (Cryo-EM) structures, suggesting that these clusters have low signal-to-noise ratio and are less informative than the remainder of the clusters.

**Figure 1.**
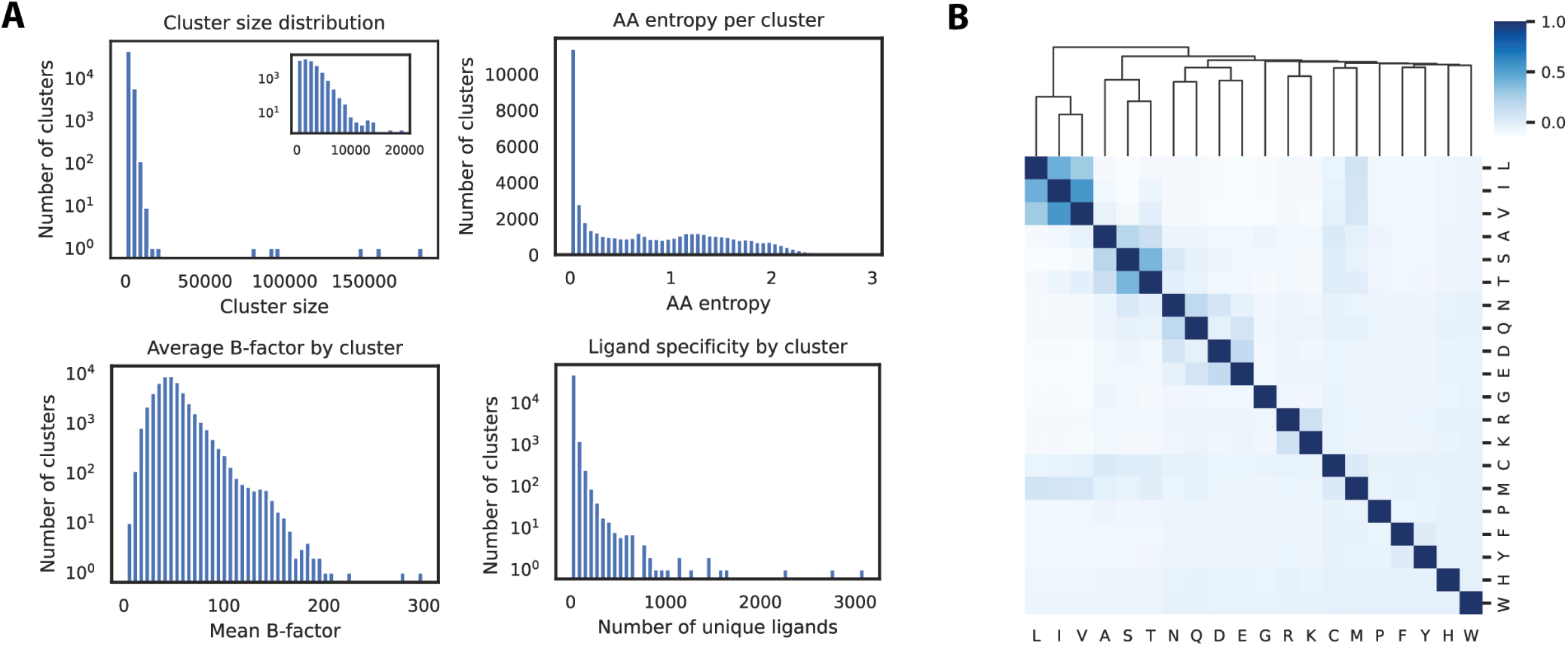
Evaluation and characterization of clusters across the full PDB database. **A.** Clockwise from top left, we show histograms of (1) cluster sizes (log scale) with inset showing only the subset of clusters with up to 20,000 environments; (2) entropy (i.e. conservation) of the central amino acid of each cluster; (3) Number of unique ligands in contact with environments in each cluster, based on PDB data processed by BioLIP; and (4) average crystallographic B-factor of the central residue over environments in each cluster. **B.** Correlation matrix for central residue frequencies over all clusters. Each square *i, j* represents the Pearson correlation between the frequencies of residue *i* and *j* over all clusters. Rows and columns are hierarchically clustered to identify patterns of residue substitutability within structural clusters.

We then characterized each cluster based on the structural and biochemical features of its constituent environments. First, we find that most clusters consist of environments centered around a diversity of residue types, with only 8.7% being associated with a single amino acid. The remaining clusters have a relatively uniform entropy distribution, confirming that the clusters capture more local structural information than the amino acid identity alone (Fig. 1A). By measuring the correlations between the distributions of each amino acid, we also assess the degree to which different residues co-occur in the same environments (Fig. 1B). This analysis reveals that the co-occurrence patterns correspond to the expected substitutability of the amino acids. For example, we observe correlations between small nonpolar amino acids leucine (L), isoleucine (I), and valine (V), negatively charged aspartic acid (D) and glutamic acid (E), and sulfur-containing cysteine (C) and methionine (M). Then, we extracted the residue-level B-factor of each residue from the corresponding PDB file as a proxy for the level of structural order in each environment. The clusters exhibit a range of average B-factor with a slight rightward skew, as shown in Fig. 1A, with a few notable outliers corresponding to clusters in highly flexible regions.

Finally, we mapped each environment to a set of external databases to provide additional insights into the potential structural and functional role of each cluster. These include protein-level annotations such as SCOP classification, Gene Ontology (GO) codes, and enzyme commission (EC) numbers as well as residue-level annotations such as the set of ligands in contact with each residue in the PDB. The ligand information enables insights into the functional interactions that each structural motif may be involved in, including peptide, nucleic acid, metal ion, and small molecule interactions. The clusters display a wide range of specificity to ligands, with some interacting with over a thousand unique ligands (Fig. 1A). Notably, this data is strongly biased towards the protein-ligand interactions that are more frequently studied and deposited in the PDB, and many clusters likely interact with ligands which have simply not been confirmed by co-crystal structures. For instance, the majority of the 3,101 unique ligands which are in contact with environments from cluster 20,672 are ATP/ADP analogs, mimetics, or competitive inhibitors, an extremely common and well-studied class of compounds.

### 3.2 Protein-level representations based on cluster frequencies

We developed a fixed-length fingerprint representation based on the frequencies of each cluster in a protein structure (see Methods 2.4). These fingerprints enable efficient comparison between protein structures and provide a simple representation for developing machine learning methods on protein-level tasks. We evaluate the capabilities of our cluster-based fingerprints for each purpose using two tasks which require that a representation captures complex aspects of a protein’s structure: structure search and decoy ranking. For all-vs-all searching in the SCOP40 database, a simple cosine similarity between fingerprints has comparable precision to Foldseek over the top *k* results for each values of *k* up to *k* = 10 (Fig. 2A).

**Figure 2.**
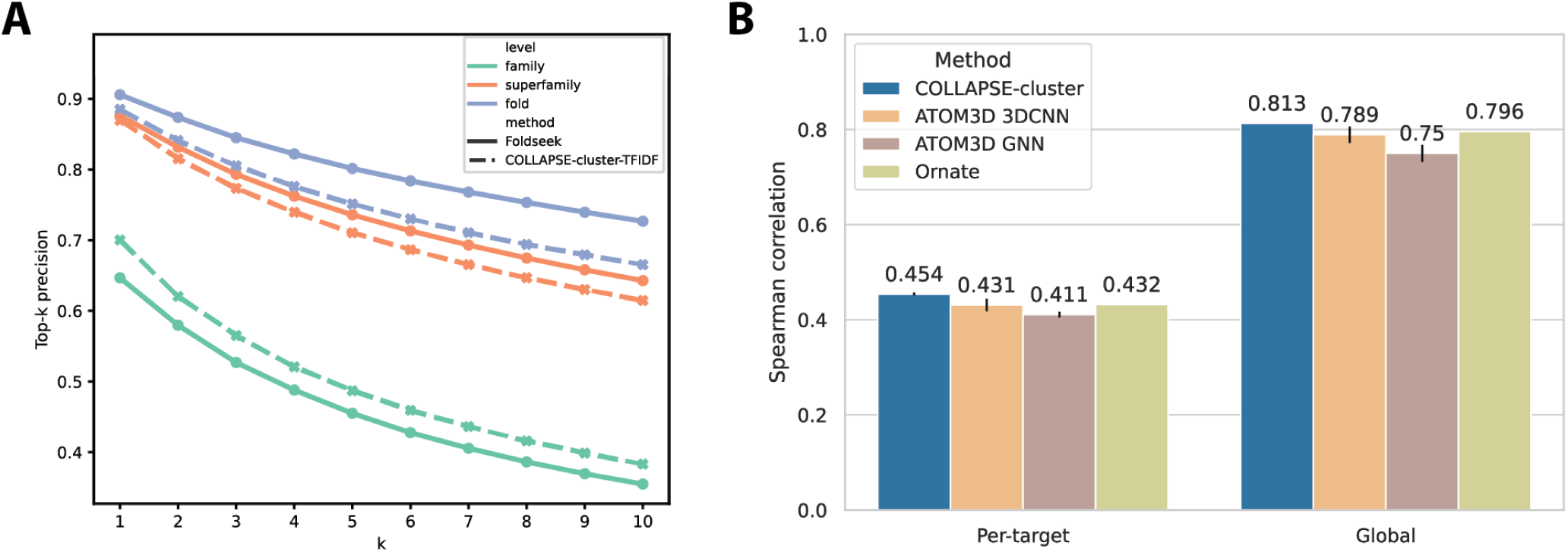
Protein-level fingerprints based on cluster frequencies. **A.** Performance of cluster-based fingerprints on all-vs-all structure similarity search in SCOP40 database. Precision of the top *k* most similar search results is shown at each level of the SCOP hierarchy, from most to least specific: family (green), superfamily (orange), and fold (blue). The results for cluster-based fingerprints are shown with dashed lines, and the Foldseek baseline is shown with solid lines. **B.** Benchmarking results for protein structure ranking (PSR) task from ATOM3D. Results for a linear predictor trained on cluster-based fingerprints are shown in blue compared to two ATOM3D baseline models—3D convolutional neural network (3DCNN) and graph neural network (GNN)—and state-of-the-art method Ornate, all trained end-to-end on the atomic protein structure. Performance metric is per-target and overall (global) Spearman correlation between predicted and actual GDT_TS score to the unseen target structure, and results for baselines are taken directly from ATOM3D^32^. Bar heights and error bars represent the mean and standard deviation over three independent training runs.

While Foldseek performs slightly better at higher levels of the SCOP hierarchy (superfamily and fold), our cluster-based fingerprints achieve higher precision for identifying matches at the most specific level (family). The top-1 performance is particularly favorable, achieving accuracy of 70.0% (vs. 64.7%) for families and very comparable accuracies of 87.3% and 89.2% (vs. 87.6%, 90.6%) for superfamilies and folds, respectively. For assessing the quality of predicted protein structures relative to their targets, a simple feed-forward predictor trained on the cluster fingerprints outperforms all baselines from the ATOM3D benchmark dataset (Fig. 2B), both in terms of average per-target Spearman correlation (0.454 *±* 0.003) and global correlation (0.813 *±* 0.000) across all targets.

### 3.3 Modeling kinase active site similarity and cross-reactivity

Cluster-based representations are not only effective for modeling whole protein structures, but they can also be used to compare important functional sites or regions within proteins. We explore the ability of our clusters to capture functionally relevant information in the context of kinase active sites. Kinases play a crucial role in cellular signaling by catalyzing the phosphorylation of serine, threonine, and tyrosine residues, and dysregulation of this signaling is implicated in cancer and a wide range of other diseases. The development of small-molecule kinase inhibitors has therefore become a major focus of pharmaceutical research, leading to several breakthrough drugs^50, 51^. However, these drugs exhibit cross-reactivity within and between kinase subfamilies due to high structural conservation within the active site, sparking interest in the development of inhibitors that have targeted selectivity to a small number of specific kinases^52, 53^. Rational development of these next-generation inhibitors requires accurate structural modeling and comparison of kinase active sites. We sought to use our cluster-based fingerprint representations to produce an inhibitor-agnostic representation of kinase binding pockets which would better capture kinase-specific structural features.

First, we assessed the ability of our fingerprints to identify family-specific features of the binding pocket. We computed a silhouette score for each kinase structure based on the distances to other kinases in the same family and the distances to each kinase in the next-closest family. For all 95 families in the KLIFS database, the average silhouette score for our cluster-based fingerprints is greater than zero, indicating good clustering of the kinase families (Fig. 3A, Fig. S2). Among the 80 families with 10 or more examples, 63 have a significantly greater silhouette score using our cluster-based fingerprints than using the KLIFS interaction fingerprints. None of the cases where the average KLIFS IFP score is greater are statistically significant. Qualitatitive visualization of the fingerprints (Fig. 3B) also supports the fact that our structural clusters enable much better separation between kinase families than the IFP representations.

**Figure 3.**
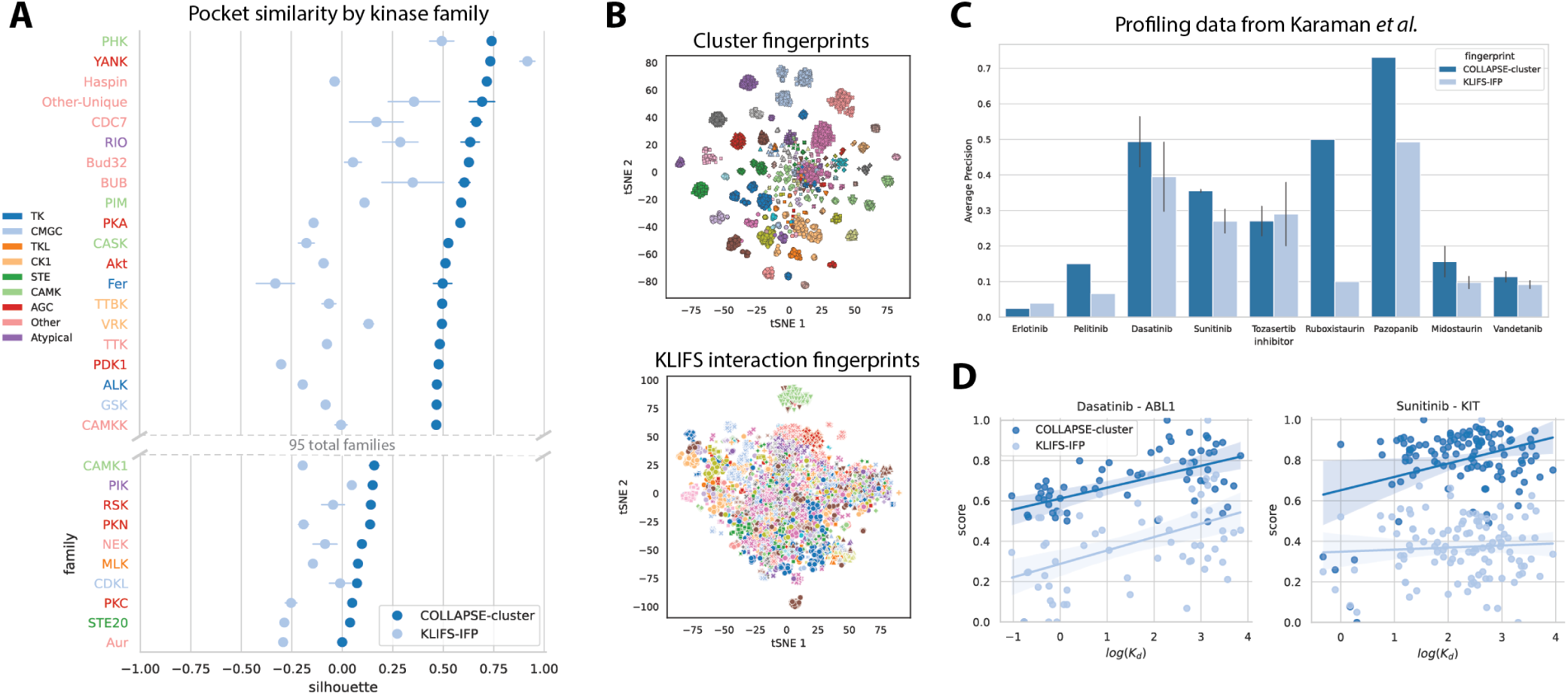
Cluster-based comparison of kinase inhibitor binding pockets. **A.** Quantitative evaluation of binding pocket similarity measured by our cluster-based fingerprints (dark blue) relative to KLIFS interaction fingerprints (light blue). Each row represents a single kinase family, and the dots represent the mean silhouette score over all kinase pockets within that family. Errorbars represent the standard error of the mean. Row labels are colored by the high-level kinase group that each family belongs to. We only show the top 20 and bottom 10 families by cluster-based silhouette score for brevity. **B.** Qualitative evaluation of binding pocket similarity by using tSNE^54^ to reduce fingerprints to two dimensions. Each dot represents a single kinase structure and is colored by the kinase family it belongs to. **C.** Identification of off-target binding interactions for nine kinase inhibitors based on experimental data from Karaman *et al.*. Bars show average precision of kinase pockets ranked by fingerprint similarity relative to a binary measure of off-target binding at 100 nM affinity. Error bars represent standard error of the mean for inhibitors with more than one kinase target. **D.** Scatterplots showing correlation between pocket fingerprint distance and quantitative binding affinity (in *log*(*K_d_*)) for two selected inhibitor-target pairs: Dasatinib–ABL1 and Sunitinib–KIT. Linear regression lines are shown along with 95% confidence intervals computed by 1000 bootstrap iterations.

Next, we investigated whether this improved modeling of the binding pocket translates to an ability to better capture the selectivity profile of kinase inhibitors. Figure 3C shows that we are able to identify off-target interactions more effectively using cluster fingerprint similarity than IFP similarity for seven of the nine kinase inhibitors tested at an affinity cutoff of 100 nM (Erlotinib and Tozasertib have comparable performance). Additionally, the similarity between pocket representations have a statistically significant correlation with the experimentally measured binding affinity for seven of 14 inhibitor-target pairs, compared to only three for IFP fingerprints (Table S2). Two such examples are shown in Fig. 3D: Dasatinib – ABL1 (*ρ_cluster_* = 0.529*, ρ_IFP_* = 0.464) and Sunitinib – KIT (*ρ_cluster_* = 0.357*, ρ_IFP_* = 0.060).

For Sunitinib in particular, the improved performance of cluster-based representations can be attributed largely to the much better identification of the few very strong binders (*log*(*K_d_*) *<* 1, bottom left) compared to the large number of weak binders in the dataset.

### 3.4 Enrichment of human disease-associated variants

We identified 13,569 clusters which were associated with at least one pathogenic mutation in the AlphaFold human proteome database, 2,056 of which had a statistically significant enrichment for pathogenic variation relative to benign variants or variants of unknown significance. Of these, 104 contain at least 5 unique variants, which we term “pathogenic clusters”. Many of these pathogenic clusters are associated with variants in cysteine residues, including Cluster 6044 (Fig. 4A; Table S3). This cluster includes 18 proteins with known missense variants, of which 82 out of 85 are annotated as pathogenic. These mutations are associated with a diverse range of diseases, including blood (e.g. hemophilia B, thrombophilia, factor VII deficiency), connective tissue (e.g. Marfan syndrome, congenital contractural arachnodactyly), malformation (e.g. Cenani-Lenz syndactyly syndrome, Adams-Oliver syndrome, Hennekam lymphangiectasia-lymphedema), and vision (e.g. retinitis pigmentosa) disorders.

**Figure 4.**
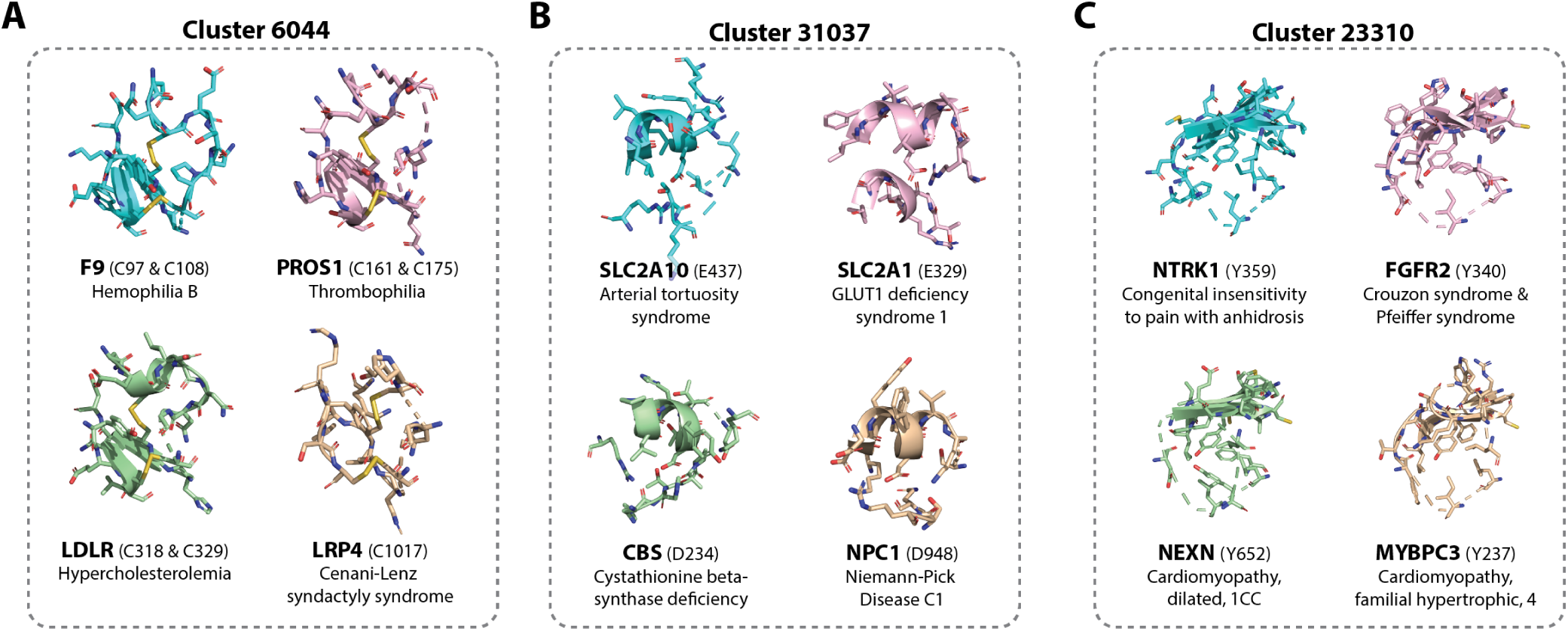
Three example clusters with statistically significant enrichment for human pathogenic variants. For each cluster we show the 5Å environments surrounding the wild type amino acid for four sampled variants, as well as the disease associated with that variant. **A.** Cluster 6044, which is characterized by mutations in a disulfide bonding motif. **B.** Cluster 31,037, which is characterized by mutations in a negatively charged central residue. **C.** Cluster 23,310, which is characterized by mutations in a central tyrosine.

Many pathogenic clusters also involve mutations in other residue types, such as Cluster 31,037, which is centered around variants which affect negatively charged aspartic acid or glutamic acid residues in a short helical motif (Fig. 4B; Table S4). This cluster contains nine proteins with 11 unique variants, all pathogenic. These variants are associated largely with rare metabolic diseases (e.g. Niemann-Pick disease, GLUT1 deficiency syndrome, isovaleric acidemia, cystathionine beta-synthase deficiency), but also with developmental disorders (Nicolaides-Baraitser syndrome, autism spectrum disorder) and others. Another 10 pathogenic variants are found in Cluster 23,310, which involves a mutant tyrosine residue in a central beta-sheet motif (Fig. 4C; Table S5). Three of these variants are associated with cardiomyopathy, but others are linked to developmental (Crouzon and Pfeiffer syndromes), neurological (congenital insensitivity to pain with anhidrosis), and endocrine (hypogonadotropic hypogonadism) disorders. Each of the clusters enriched for pathogenic variation in humans is associated with a strongly conserved central residue surrounded by a relatively tightly packed local structural environment, reflecting the fact that these types of structural motifs are less tolerant of missense variation.

In total, 44% (9,703 out of 21,834) of human proteins in the AlphaFold structure database contain at least one pathogenic cluster, suggesting that previously unobserved mutations in any of the corresponding residues in these proteins would result in high risk of pathogenicity. All data for these potentially deleterious residue locations are published in Supplementary Dataset 1 as a resource to help in evaluating novel variants.

## 4 Discussion

In this work, we perform unsupervised clustering of protein sites to discover the landscape of conserved local building blocks that make up protein structures. By learning the clusters over 15 million sites and characterizing them on over 100 million sites across the entire PDB, this makes up the largest-scale structural analysis of protein sites to date. We were able to achieve this due to the use of general-purpose pre-trained COLLAPSE embeddings which reduce the complex 3D geometry of a protein site to a continuous embedding. The high level of structural conservation within these clusters, even when they correspond to different amino acids and belong to proteins with disparate global folds, is evidence that the embeddings capture local atomistic geometry with high fidelity.

Based on intrinsic measures of cluster compactness and separation (WCSS and silhouette scores), we observed that there was not a single value of *k* which clearly results in optimal clustering. This suggests that local protein structure does not separate into a single discrete set of conformations but rather occupies a continuum of variation, much like the relationship between folds at the domain level observed by previous works. Therefore, the choice of the number of clusters, *k*, is a trade-off between the specificity of each cluster and its utility for cross-protein analysis (i.e. in the limit where each cluster is present in only one protein, the clusters are no more useful than assigning all residues with a unique label). Additionally, as *k* increases, the training and downstream application of the clusters becomes computationally inefficient. We find empirically that 50,000 clusters provide a good balance of these factors. We also note that this is similar in order of magnitude to the number of discrete input categories present in other well-established biomedical datatypes (e.g. the number of words in the English language or the number of genes in the human genome), so we posit that this choice provides a strong basis for many established computational approaches to be adapted for protein structure analysis.

To illustrate this, we devise a fixed-length protein representation based on the normalized frequencies of each cluster in the protein structure. This approach takes inspiration from the term frequency-inverse document frequency (TF-IDF) representation of documents used in text mining and information retrieval, and is based on the intuition that a protein’s global structure and function are determined by its specific combination of local sites. We show through our experiments that this approach is sufficient to identify proteins from the same SCOP family with greater precision than Foldseek, the well-established leader in structure-based search. The improvement over Foldseek at the most specific level of comparison may be due to the fact that COLLAPSE embeddings are based on all-atom representations of each environment (including side chains) while Foldseek’s structure alphabet accounts for only backbone conformation. Additionally, our cluster-based representations enable state-of-the-art performance on the task of ranking the accuracy of predicted protein structures relative to an unseen target. This is a difficult task that relies on the ability of a model to identify low-energy conformations which are closest to the native structure. The high performance of our fingerprints with only a simple linear prediction head suggests that the frequency of clusters alone is enough to identify “native-like” conformations at least as accurately as complex models trained end-to-end on the structures themselves. This result has implications for the utility of site-based approaches for evaluating not only computationally predicted structures but also the feasibility of *de novo* designed proteins. In general, the success of this simple “bag-of-sites” approach is a promising sign that combining site-level structural clusters with spatial or sequential information could enable even more powerful protein structure representations.

A major benefit of using a site-based approach to protein representation is that it is easy to perform analysis at multiple scales of protein structure. Many functional regions in proteins, such as ligand binding pockets, are larger than the environment around a single amino acid but more localized than the entire protein chain. We show that cluster-based representations of kinase active sites can be used to effectively separate kinase families, showing that the sites contain useful functional information. Additionally, we show that the similarity between cluster-based representations can be used to identify off-target effects more accurately than interaction fingerprints which explicitly account for ligand-protein interactions. This is important because our method is ligand-agnostic and can be used to evaluate any pair of binding pockets regardless of whether they have known co-crystal structures, making it a powerful tool for evaluating the selectivity of new kinase inhibitors. Expanding this analysis beyond kinases to identify binding pocket similarities across different protein families would also be useful for identifying novel drug off-target interactions and potential repurposing opportunities.

Mapping each environment to a discrete cluster also enables us to associate structural motifs with a variety of information such as amino acid conservation, structural disorder, and ligand interactions. This type of metadata is useful for evaluating the functional importance of individual sites within a protein, and detecting statistical associations within this data can elucidate previously unknown structure-function relationships. Additionally, the AlphaFold structure database provides the opportunity to discover such patterns over entire proteomes without being limited to experimentally solved structures. Our analysis of genetic variation in the human proteome illustrates this by identifying structural motifs associated with mutation pathogenicity. Understanding the structural environment surrounding a mutated amino acid can be important for improving prediction of pathogenicity for variants of unknown function and for elucidating the molecular mechanism underlying a variant’s clinical effect (e.g. by affecting protein stability or by impacting the biochemistry of a functional site). In other cases, identifying sites which can tolerate mutations and are functionally important can also be useful for designing new protein variants with improved fucntional characteristics^55^. In addition to the 104 clusters we identified to be associated with pathogenicity, we also found many clusters which were not associated with enough variants to achieve statistical significance, suggesting that including more variant data would yield improved results. This could be done by expanding data to proteins from other organisms or variant annotations from other sources. For example, AlphaMissense^34^ recently released predicted variant annotations for the entire human proteome, which could be used to supplement known clinical annotations and improve power. The known challenges of computationally predicting individual variant effects (AlphaMissense only modestly improves over previous methods and still has low correlation with experimental data^56, 57^) may also be mitigated by considering many variants within each cluster.

In conclusion, we present a large-scale characterization of the landscape of conserved structural motifs in proteins and provide evidence that this lexicon of motifs enables new approaches to modeling proteins from a site-based perspective. We believe that a site-centric approach is important for understanding functional relationships as protein structure databases continues to expand. Additionally, as deep learning produces new breakthroughs in structure-based protein engineering and design, it is becoming increasingly important to accurately represent structure-function relationships at amino acid resolution in order to produce proteins with specific functional characteristics. We anticipate that the structural clusters identified by this work and the corresponding database will provide a foundation for improved site-based modeling of protein structures and enable the discovery of new functional relationships across the protein universe.

## Acknowledgments

Computing for this project was performed on the Sherlock cluster; we would like to thank Stanford University and the Stanford Research Computing Center for providing computational resources and support. This work is supported by Chan-Zuckerberg Biohub and the National Institutes of Health (GM102365 and LM012409).

## Supplementary materials

**Figure S1.**
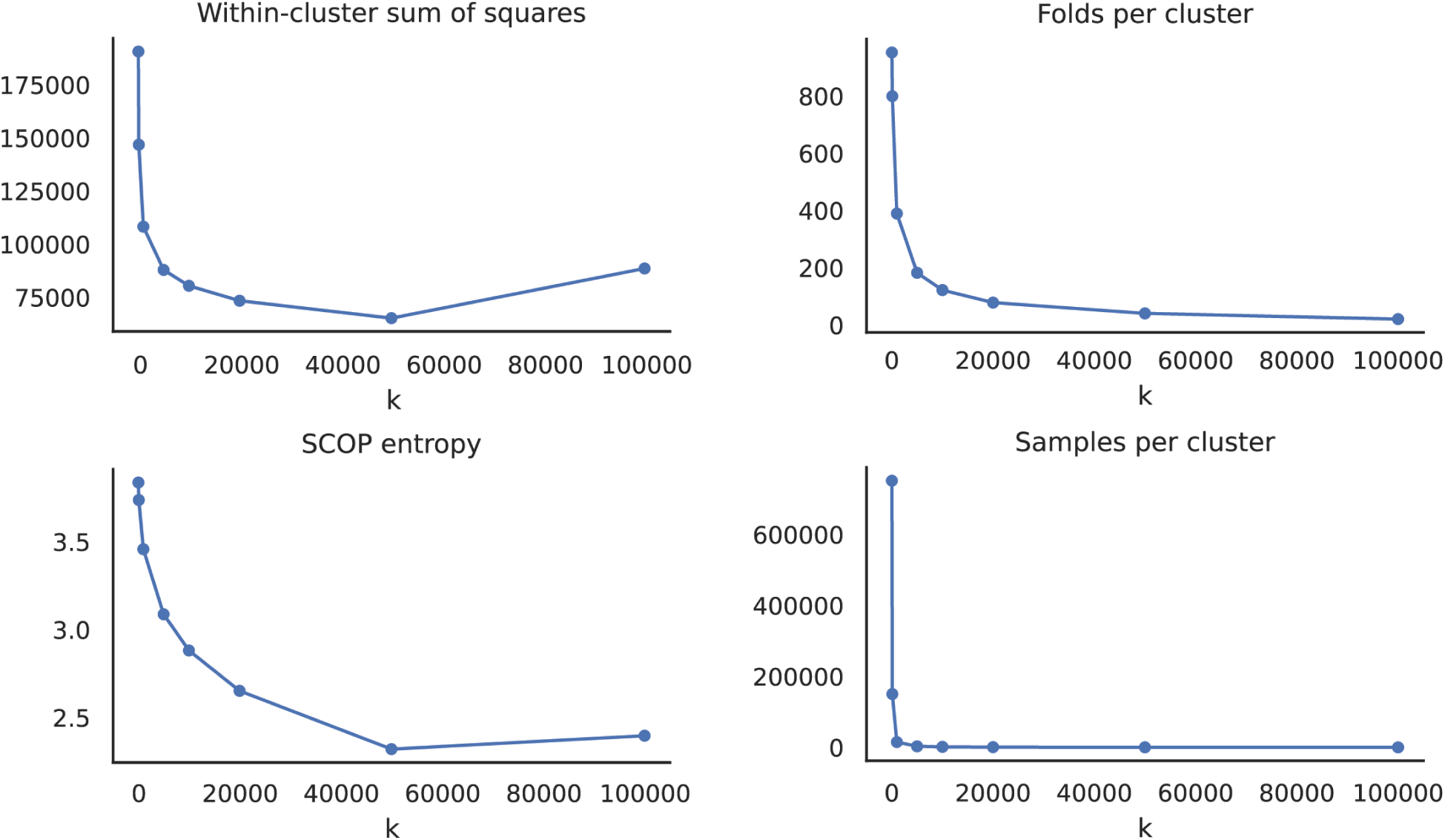
Cluster characteristics at varying *k*. Clockwise from top left: (1) within-cluster sum of squares, (2) unique SCOP folds per cluster, (3) entropy of SCOP families within each cluster, and (4) average number of environments in each cluster.

**Figure S2.**
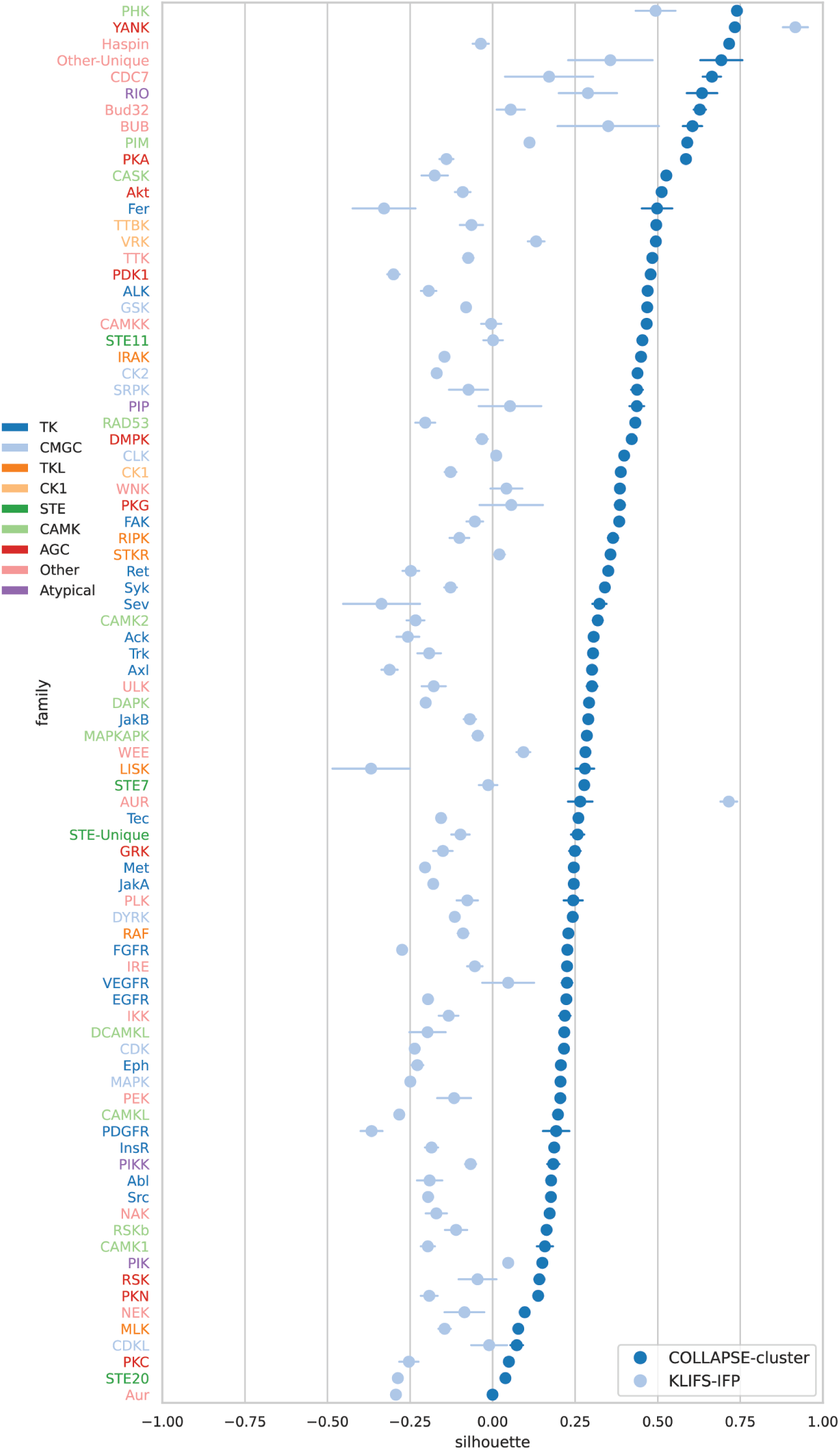
Silhouette scores of pocket similarity for all 95 kinase families. Each row represents a single kinase family, and the dots represent the mean silhouette score over all kinase pockets within that family. Errorbars represent the standard error of the mean. Row labels are colored by the high-level kinase group that each family belongs to.

**Table S1.** Kinase inhibitors evaluated in this study, as well as their primary target(s) as specified by Karaman *et al.*^46^.

| Inhibitor name | Alternative name<br>(Karaman *et al.*) | Primary target(s) |
| --- | --- | --- |
| Erlotinib |  | EGFR |
| Pelitinib | EKB-569 | EGFR |
| Vandetanib | ZD-6474 | EGFR, RET |
| Dasatinib |  | ABL1, SRC |
| Sunitinib |  | KIT, FLT3 |
| Pazopanib | GW-786034 | FLT1, FLT4 |
| Midostaurin | PKC-412 | FLT3, KIT |
| Ruboxistaurin | LY-333531 | PKCb |
| Tozasertib | VX-680/MK-0457 | AurA, AurB, AurC |

**Table S2.** Spearman rank correlation between fingerprint similarity and experimental binding affinity for each kinase inhibitor-target pair. Bold indicates the best correlation for each pair, and stars represent Bonferroni-corrected statistical significance: *** (*p <* 0.01), ** (*p <* 0.05), * (*p <* 0.1).

| Inhibitor | Target | COLLAPSE-cluster | KLIFS-IFP |
| --- | --- | --- | --- |
| Dasatinib | ABL1 | **0.529\*\*\*** | 0.464*** |
|  | SRC | **0.533\*\*\*** | 0.289 |
| Erlotinib | EGFR | 0.141 | **0.356** |
| Pelitinib | EGFR | -0.040 | **0.180** |
| Midostaurin | FLT3 | 0.013 | **0.083** |
|  | KIT | **0.092** | -0.084 |
| Sunitinib | FLT3 | **0.348\*\*\*** | 0.177 |
|  | KIT | **0.357\*\*\*** | 0.060 |
| Tozasertib | AurA | **0.215** | -0.018 |
|  | AurB | 0.332** | **0.337\*\*** |
|  | AurC | **0.327\*\*** | 0.280 |
| Vandatenib | EGFR | 0.148 | **0.168** |
| Pazopanib | FLT1 | **0.659\*\*\*** | 0.460** |
| Ruboxistaurin | PKCb | 0.250 | **0.284** |

**Table S3.** Protein metadata for all mutations in Humsavar associated with cluster 6044, organized by Uniprot ID. Asterisks (*) indicate non-pathogenic mutations.

| Uniprot ID | Protein name | Gene name | Mutations | Diseases |
| --- | --- | --- | --- | --- |
| O75096 | Low-density lipoprotein receptor-related protein 4 | LRP4 | C1017R | Cenani-Lenz syndactyly syndrome (CLSS) |
| P00740 | Coagulation factor IX | F9 | C97S, C108S, C134Y | Hemophilia B (HEMB) |
| P01130 | Low-density lipoprotein receptor | LDLR | C318Y*, C318F, C318R, C329Y, C329F, C358Y, C364R*, C368R, C368Y* | Hypercholesterolemia, familial, 1 (FHCL1) |
| P07225 | Vitamin K-dependent protein S | PROS1 | C121Y, C161G, C175F, C247G | Thrombophilia due to protein S deficiency, autosomal dominant (THPH5) |
| P07911 | Uromodulin | UMOD | C77Y, C126R, C170Y, C300G, C315R | Tubulointerstitial kidney disease, autosomal dominant, 1 (ADTKD1) |
| P08709 | Coagulation factor VII | F7 | C121F, C151S | Factor VII deficiency (FA7D) |
| P23352 | Anosmin-1 | ANOS1 | C163Y, C163R, C172R | Hypogonadotropic hypogonadism 1 with or without anosmia (HH1) |
| P24043 | Laminin subunit alpha-2 | LAMA2 | C527Y | Merosin-deficient congenital muscular dystrophy 1A (MDC1A) |
| P35555 | Fibrillin-1 | FBN1 | C504F, C504R, C546W, C576Y, C587Y, C727Y, C781R, C781Y, C811Y, C926R, ..., [46 more] | Marfan syndrome (MDS); Ectopia lentis 1, isolated, autosomal dominant (ECTOL1) |
| P35556 | Fibrillin-2 | FBN2 | C1246F, C1257W, C1257R, C1384F, C1384Y, C1253Y, C1253W, C1257W, C1257R, C1268R, C1253Y, C1253W | Contractural arachnodactyly, congenital (CCA) |
| P46531 | Neurogenic locus notch homolog protein 1 | NOTCH1 | C429R | Adams-Oliver syndrome 5 (AOS5) |
| P78504 | Protein jagged-1 | JAG1 | C911Y | Alagille syndrome 1 (ALGS1) |
| P82279 | Protein crumbs homolog 1 | CRB1 | C310Y, C891G, C939Y, C1181R, C1223S | Retinitis pigmentosa 12 (RP12); Leber congenital amaurosis 8 (LCA8); |
| Q5IJ48 | Protein crumbs homolog 2 | CRB2 | C620S | Focal segmental glomerulosclerosis 9 (FSGS9) |
| Q5T1H1 | Protein eyes shut homolog | EYS | C1176R | Retinitis pigmentosa 25 (RP25) |
| Q6UXH8 | Collagen and calcium-binding EGF domain-containing protein 1 | CCBE1 | C102S | Hennekam lymphangiectasia-lymphedema syndrome 1 (HKLLS1) |
| Q92832 | Protein kinase C-binding protein NELL1 | NELL1 | C553F* | N/A |
| Q9HC23 | Prokineticin-2 | PROK2 | C46Y | Hypogonadotropic hypogonadism 4 with or without anosmia (HH4) |

**Table S4.** Protein metadata for all mutations in Humsavar associated with cluster 31037, organized by Uniprot ID. Asterisks (*) indicate non-pathogenic mutations.

| Uniprot ID | Protein name | Gene name | Mutations | Diseases |
| --- | --- | --- | --- | --- |
| O15118 | NPC intracellular cholesterol transporter 1 | NPC1 | D948N, D948H, D948Y | Niemann-Pick Disease C1 (NPC1) |
| O95528 | Solute carrier family 2, facilitated glucose transporter member 10 | SLC2A10 | E437K | Arterial tortuosity syndrome (ATORS) |
| P11166 | Solute carrier family 2, facilitated glucose transporter member 1 | SLC2A1 | E329Q | GLUT1 deficiency syndrome 1 (GLUT1DS1) |
| P26440 | Isovaleryl-CoA dehydrogenase, mitochondrial | IVD | D72N | Isovaleric acidemia (IVA) |
| P35520 | Cystathionine beta-synthase | CBS | D234N | Cystathionine beta-synthase deficiency (CBSD) |
| P51531 | Probable global transcription activator SNF2L2 | SMARCA2 | D1158V | Nicolaides-Baraitser syndrome (NCBRS) |
| Q7Z3K3 | Pogo transposable element with ZNF domain | POGZ | E1040K | Autism spectrum disorder (ASD) |
| Q9NRW7 | Vacuolar protein sorting-associated protein 45 | VPS45 | E238K | Neutropenia, severe congenital 5, autosomal recessive (SCN5) |
| Q9Y5Z9 | UbiA prenyltransferase domain-containing protein 1 | UBIAD1 | D236E | Corneal dystrophy, Schnyder type (SCCD) |

**Table S5.**
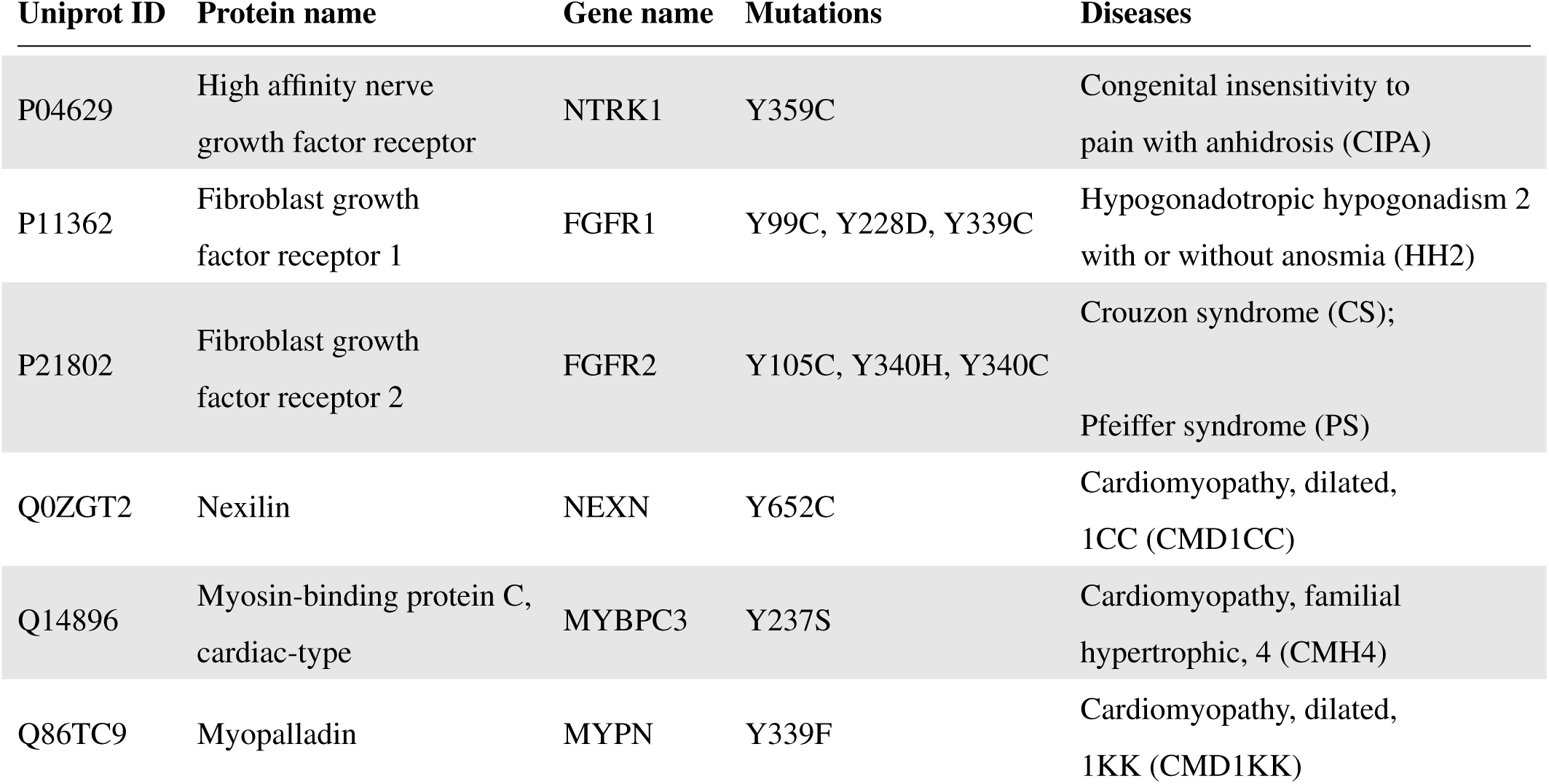
Protein metadata for all mutations in Humsavar associated with cluster 23310, organized by Uniprot ID. Asterisks (*) indicate non-pathogenic mutations.

